# Natural killer cell division regulates FcεRIγ expression downstream of mTOR activity

**DOI:** 10.1101/2021.08.03.454985

**Authors:** Avishai Shemesh, Daniel R. Calabrese, Janice Arakawa-Hoyt, John R. Greenland, Lewis L. Lanier

**Affiliations:** Department of Microbiology and Immunology, University of California, San Francisco, San Francisco, California 94143, USA; Parker Institute for Cancer Immunotherapy, University of California, San Francisco, San Francisco, CA 94143, USA; Department of Medicine, University of California, San Francisco, CA; Medical Service, Veterans Affairs Health Care System, San Francisco, CA

## Abstract

The expansion of human FcεRIγ^-/low^ (FcRγ^-/low^) natural killer (NK) cells accrues during viral infections; however, the molecular mechanisms regulating FcRγ expression is not well defined and can have implication for host protection and NK cell immunotherapy. Our analysis of NK cell subsets in lung transplant patients during rapamycin treatment revealed significantly lower FcRγ levels in the NK cell population. Moreover, lower FcRγ levels in healthy donors were associated with low mTORC1/C2 activity and low T-bet expression. Cell division suppression by rapamycin or TGFβ suppressed FcRγ upregulation during IL-2 receptor stimulation, whereas promoting NK cell division by co-inhibiting FOXO1 activity restored FcRγ upregulation. These results suggest that the human FcRγ^-/low^ NK cell phenotype is associated with cell division suppression and reduced mTOR activity.

## Introduction

NK cells are innate lymphocytes functionally regulated by activating and inhibitory germline-encoded receptors^1^. The major developmental stages of human NK cells are defined by CD16 and the density of CD56 expression. Immature NK cells in the peripheral blood are CD3^-^CD56^bright^CD16^-^, whereas mature NK cells are CD3^-^CD56^dim/-^CD16^+ 2^. Downregulation of the inhibitory receptor NKG2A and upregulation of Killer-cell Immunoglobulin-like Receptors (KIRs) and the senescent cell marker CD57 further accompany human CD56^dim^CD16^+^ NK cell maturation ^3,4^.

In NK cells, the adaptor protein FcεRIγ (FcRγ) is expressed as a homodimer or heterodimer with the adaptor protein CD3ζ to regulates the expression and function of the activating receptors CD16, NKp30, and NKp46^1,5^. In humans, immature CD56^bright^CD16^-^ NK cells express lower FcRγ levels, whereas FcRγ levels increase with the development of mature CD56^dim^CD16^+^FcRγ^+^ NK cells. Human cytomegalovirus (HCMV) infection can lead to the expansion of a subset of mature CD56^dim^CD16^+^CD57^+/-^FcRγ^-/low^ NK cells that exhibit stable population frequencies over years ^6,7^. Reduced FcRγ levels in mature CD56^dim^CD16^+^ can be associated with lower surface NKp30 and NKp46 levels, which may lead to reduced NK cell responses against tumors expressing ligands for these activating receptors ^8–11^, yet have a minor influence on CD16 expression in vivo ^5^. Moreover, CD56^dim^CD16^+^FcRγ^-/low^ NK cells exhibit a higher CD16-mediated response, possibly due to its exclusive association with the CD3ζ homodimer or the CD57 phenotype ^4,12,13^. In some individuals, a unique subset of mature CD56^dim^CD16^+^NKG2A^-^NKG2C^high^ NK cells are expanded during acute HCMV infection, leading to their designation as “adaptive” NK cells ^14,15^. However, NKG2C-negative CD56^dim^CD16^+^NKG2A^+^FcRγ^-/low^ or CD56^dim^ CD16^+^NKG2A^-^FcRγ^-/low^ NK cells are also present both in HCMV-infected and uninfected individuals, indicating they may be expanded due to stimuli other than HCMV ^16,17^. In addition, in some individuals, lower FcRγ levels are associated with genes such as *SYK, SH2D1B* (EAT2), and/or *ZBTB16* (PLZF) which are downregulated in adaptive CD56^dim^FcRγ^-/low^ NK cells by DNA methylation ^11,12,18^. FcRγ protein loss is reported during CD16 stimulation of human NK cells, while IL-2 restored FcRγ expression indicating FcRγ expression is regulated by the IL-2 receptor signaling ^19^, yet to date, the molecular mechanism regulating FcRγ expression in human NK cells downstream of IL-2 receptor is not well characterized and might affect NK cell-mediated immune response in organ transplantation, viral infection, or several malignancies^20^.

The IL-2 receptor is a heterotrimeric complex that signals through CD122 (IL-2Rβ) and CD132 (IL-2Rγ) to regulate NK cell homeostasis, proliferation, and differentiation by interacting with IL-2 or IL-15 as soluble or membrane-bound forms^21–24^. IL-2Rβγ binds to soluble IL-2 or IL-15 with intermediate affinity (*K*_d_ ≈ 10^−9^ M). Higher binding affinity of IL-2 (*K*_d_ ≈ 10^−11^ M) is mediated by IL-2 cis-presentation via IL-2Rα (CD25), which is *de novo* translated in NK cells during cell activation leading to the formation of a heterotrimeric IL-2Rαβγ complex ^25^. IL-15 forms a heterotrimeric receptor complex with a similar high-affinity binding to IL-15Rα (CD215) trans-presented by myeloid cells ^23^. Both IL-2 and IL-15 impact NK cell expansion during MCMV infection^26^. IL-2Rβγ signaling leads to the phosphorylation of STAT5 (pSTAT5) and STAT4 (pSTAT4), and other STATs directly or indirectly, which are essential for NK cell survival, priming, and cell division regulation ^27–29^

NK cell division accrues downstream of the IL-2 receptor and it’s induce via PI3K activation, leading to the phosphorylation of the kinase AKT at position T308 (pAKT^T308^) and the activation of the mechanistic targets of rapamycin complex-1 (mTORC1)^30,31^. mTORC1 activation leads to the phosphorylation of ribosomal protein S6 at the initial position S235/236 (pS6^S235/236^) and regulates cap-dependent protein translation and cell cycle progression. The mTOR protein is also part of the mTOR complex-2 (mTORC2), which phosphorylates AKT at position S473 (pAKT^S473^) and regulates FOXO1 activity ^32,33^. Inhibition of mTORC1/C2, directly or indirectly, by TGFβ or rapamycin antagonizes NK cell division ^34–37^. Impaired mTOR activity in mouse NK cells decreases CD122 and CD132 expression ^21,38,39^. Additionally, the transcription factors EOMES and T-bet, which are associated with NK cell maturation and regulated downstream of the mTOR/FOXO1 axis, are suggested to regulate CD122 expression ^40–43^.

Here we characterized the NK cell subsets in lung transplant patients before and during rapamycin treatment and found reduced FcRγ expression in all defined NK cell subsets. Additionally, we characterized STATs signaling and mTOR activity in resting (steady-state) immature and mature human NK cell subsets from healthy donors with high frequencies of NKG2C^high^ cells and found a significant association between FcRγ expression and mTOR activity and T-bet expression. *In vitro*, we showed that rapamycin or TGFβ suppressed FcRγ upregulation during IL-2 stimulation, which was significantly correlated with NK cell division and was salvaged by FOXO1 inhibition. These results indicate that cell division progression, downstream of mTOR activity, regulates FcRγ expression.

## Results

### Rapamycin treatment is associated with reduced FcRγ levels in NK cells from lung transplant patients

NK cells are reported to play a role in graft-versus-host disease (GvHD) associated with organ rejection or tolerance ^44–46^. Rapamycin is an immunosuppressive agent used during solid organ transplantation ^47,48^. Further, mTOR activity regulates NK cell maturation^21,38,39^. Therefore, we assessed the influence of rapamycin treatment on the frequencies of NK cell subsets by comparing matched PBMC samples obtained from four lung transplant patients before and during rapamycin treatment (Fig. 1A). Additionality we assessed samples from five lung transplant patients that were at least 1-year post-transplant and matched for transplant indication. We gated on live CD45^+^dump^-^ (CD3^-^CD19^-^CD14^-^CD123^-^CD4^-^) NKp46^+^, CD56^+^ NK cells, and sub-gated the NK cells to four NK cell subsets based on NKG2A and CD16 expression; 1. NKG2A^+^CD16^-^, 2. NKG2A^+^CD16^low^, 3. NKG2A^+^CD16^+^, and 4. NKG2A^-^CD16^+^ (Fig. 1A). NK cells percentages significantly increased during rapamycin treatment (Fig, 1A, 1B). No significant changes were detected in the frequencies of the NK cell subsets, while the percentages of CD57^+^NKG2A^-^CD16^+^ NK cells significantly increased (Fig. 1C, 1D). NKG2C expression was not associated with a defined NK cell subset or rapamycin treatment (Fig. 1E). However, all the defined NK cell subsets displayed significantly lower FcRγ levels during treatment (Fig.1F and S1), which was accompanied by a lower percentage of Ki67^+^ cells and relative pS6^S235/236+^ cells (Fig. 1G, 1H). The results suggest lower FcRγ expression is associated with immunosuppression, reduced mTOR activity and/or NK cell division.

**Figure 1:**
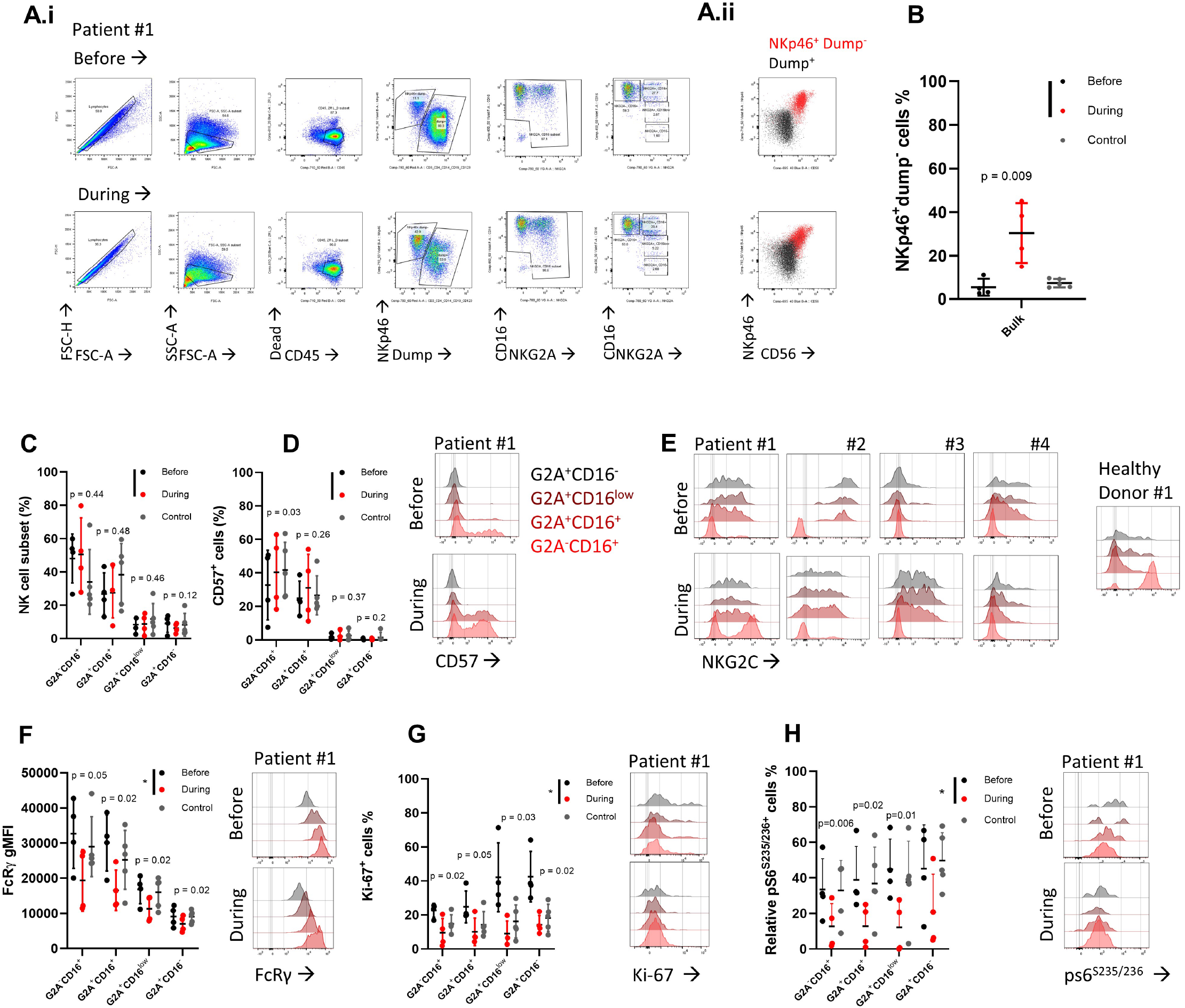
FcRγ levels in NK cells from lung transplant patients during rapamycin treatment. PBMC samples from lung transplant recipients before or during rapamycin administration (n= 4). The control donor group is subjects that were at least 1-year post-transplant and matched for transplant indication (n= 5). Mean± S.D, paired t-test, one tail, *p<0.05. **<u>(A)</u>** i. Representative flow cytometry gating analysis of matched samples from lung transplant recipients before (upper dot plots) or during (lower dot plots) rapamycin treatment, ii. Representative dot plots of NK.p46 vs. CD56 expression between NK.p46^+^dump^-^(red) and Dump^+^ cells (black) before (upper dot plot) or during (lower dot plot) rapamycin treatment. **<u>(B-C)</u>** Percentages of NKp46^+^dunp^-^NK cells (8), percentages of NK cell subsets (C). **<u>(D)</u>** Percentages of CDS^+^ cells, right: representative histograms of CD57 staining in the defined NK cell subsets (color-coded) before and during rapamycin treatment. **<u>(E)</u>** Histograms ofNKG2C expression in the defined NK cell subsets (color-coded). **<u>(F-H)</u>** FcRγ gMFI levels (F), and the percentages of Ki-67+ cells (G), and Relative pS6+ cells (H), right: representative histograms before and during rapamycin treatment in the defined NK cell subsets (color-coded).

### FcRγ expression at steady-state is associated with mTOR activity

To address if FcRγ levels are associated with mTOR activity, we examined STATs signaling and mTORC1/C2 activity at steady-state (resting) in healthy donors with an expanded CD56^dim^CD16^+^FcRγ^- or low^ NK cell population. NK cells were isolated by negative selection from PBMC to avoid indirect influence on NK cell signaling. Donor’s #1 mature CD56^dim^CD16^+^ NK cell subsets were predominantly defined as NKG2A^+^NKG2C^-^FcRγ^high^, NKG2A^-^ NKG2C^high^FcRγ^low^, and NKG2A^-^NKG2C^high^FcRγ^-^, whereas donor’s #2 NKG2A^-^NKG2C^high^ cell population was defined by FcRγ^-^, FcRγ^low^, or FcRγ^high^ subsets (Fig. 2A, S2A). Self-reactive KIR expression was mostly detected in donor #1 cells, whereas CD85J was associated with NKG2C expression in both donors (Fig. 2B)^16,49^. Further, higher frequencies of CD57^+^ cells were associated with low or absent FcRγ expression (Fig. 2C)^15^. Therefore, we compared the relative levels of expression of cell signaling molecules between CD56^dim^CD16^+^ NK cell subsets, further classified into the following subsets: 1. NKG2C^-^FcRγ^high^, 2. NKG2C^high^FcRγ^low^, and 3. NKG2C^high^FcRγ^-^, and between CD57^-^ and CD57^+^ cells within each of the defined subsets (Fig. 2D). Additionally, we compared cell signaling markers in immature CD56^bright^CD16^-^ NK cells relative to mature CD56^dim^CD16^+^ NK cell subsets as lower FcRγ levels were reported in the immature subset^7^. To assess the relative signaling level between the subsets and donors, we normalized NK subsets protein expression levels indexed to the amount of the protein in mature CD56^dim^CD16^+^NKG2C^-^FcRγ^+^CD57^-^ NK cells (index = 1).

**Figure 2:**
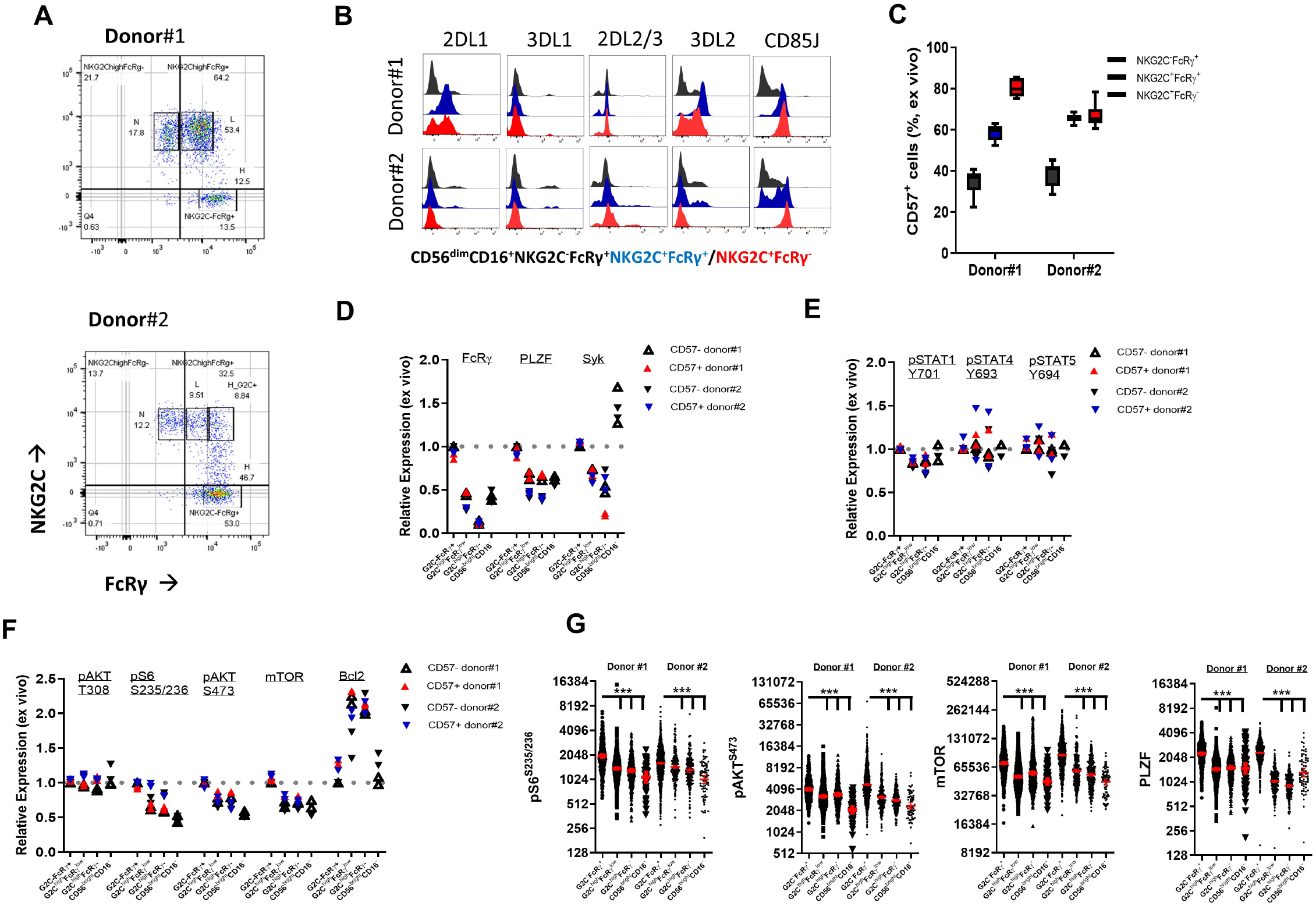
FcRγ expression at steady-state is associated with mTOR activity. **(A)** NKG2C vs. FcRγ in *ex vivo* CD56^dim^CD 16^+^ NK cells. Donor# I (upper), Donor #2 (lower). N = FcRγ^negative^, L = FcRγ^low^, H = FcRγ^high^ NK cells. **(B)** Representative histograms of indicated KIRs or CD85J on *ex vivo* CD56^dim^CD16^+^ NK cell subsets (X-axis log_10_ fluorescence). **(C)** Percentage of CDS^+^ cells in *ex vivo* CD56^dim^CD16^+^ NK cell subsets. Box and whiskers: min to max. Integrated results from 3 independent measurements. **(D-F)** Intracellular levels for informative adaptive NK cell proteins FcRγ, PLZF, Syk (D), STATs signaling molecules (pSTATl^Y701^, pSTAT4^Y963^,pSTAT5^Y964^ (E), pAKT^T308^, mTORCl/mTORC2 activity (pS6^S235/236^ and pAKT^S473^ respectively), mTOR, or Bcl2 (F), in the defined NK cell subsets. Values were normalized to CD56^dim^CD16^+^ NKG2C-FcRγ^+^CD57^-^ NK cells (index = 1). Each measurement represents independent staining of an independent blood sample collected at an independent time point. **(G)** Single cell expression of (left to right) pS6^S2351236^,pAKT^S473,^ mTOR, or PLZF, log 2 values. Red bar = geometric mean with 95% confidence interval, unpaired t-test, two tails, *p<0.05, *** p<0.001.

In line with previous reports, immature CD56^bright^CD16^-^ NK cells expressed lower levels of FcRγ similar to CD56^dim^NKG2C^high^FcRγ^low^. PLZF, but not Syk, was reduced in peripheral blood immature CD56^bright^CD16^-^ and mature CD56^dim^NKG2C^high^ NK cells (Fig. 2D). Additionally, the mature FcRγ^-^ or FcRγ^low^ NK cells expressed lower surface NKp30 and NKp46 levels, suggesting these are adaptive NK cells (Fig. S2B)^12,18^. Examination of STATs activation showed pSTAT1^Y701^ levels were lower in the NKG2C^high^FcRγ^low/-^ subsets but not in the immature CD56^bright^CD16^-^ subset, whereas no differences were detected in pSTAT4^Y693^ or pSTAT5^Y694^ (Fig. 2D). Moreover, no difference was seen in pAKT^T308^ levels. However, mTORC1 and mTORC2 activity measured by the levels of pS6^S235/236^ or pAKT^S473^, respectively, were significantly lower in adaptive NKG2C^high^FcRγ^low/-^ and immature CD56^bright^CD16^-^ NK cells. Additionally, mTOR protein levels, but not Bcl2, were significantly lower, matching FcRγ expression (Fig. 2F, 2G)^50^. CD57 expression was not associated with the reduced mTOR signaling, whereas repeated measurements on samples collected at different time points indicated these variations in cell signaling were stable (each replicate = one measurement). Examination of factors reported to regulate mTOR activity, such as pp38 MAPK^Thr180/Tyr182^, pRptor^S792^, intracellular LAMP1 (CD107a) or LAMP2 (CD107b), Rptor, and Rictor levels did not detect stable variations in these factors (Fig. S2C)^51–54^. Additionally, surface expression of the amino acid transporter subunits CD98 light chain (LAT1/SLC7A5) and CD98 heavy chain (SLC3A2) and GLUT1 were negative on all subsets of resting NK cells (data not shown)^55^. Therefore, we concluded that FcRγ expression levels *ex vivo* at steady-state are associated with low PLZF, low mTOR protein level, and low mTOR activity, but not with pSTAT1^Y701^, pSTAT4^Y693^, or pSTAT5^Y694^ levels.

### T-bet but not CD122 levels are associated with FcRγ expression

Impaired mTOR activity in mouse NK cells reduces expression of the IL-2 receptor subunits, CD122 and CD132, whereas CD57 expression in human NK cells is reported to correlate with lower CD122 levels ^3,38^ Moreover, IL-2 receptor stimulation activates mTOR and was shown to upregulate FcRγ; therefore, changes in IL-2 receptor expression might account for changes in FcRγ levels ^19,21^. Consequently, we examined the expression of IL-2 receptor subunits in the predefined resting NK cell subsets relative to FcRγ expression (Fig 2A). CD25 was absent in all NK cell subsets (data not shown). In mature CD56^dim^CD16^+^ NK cells, lower CD122 expression was associated with low or negative FcRγ expression (Fig 3A). However, CD122 levels did not couple to lower FcRγ expression in immature CD56^bright^CD16^-^ NK cells. In contrast, higher CD132 levels were associated with lower FcRγ levels in both CD56^dim^NKG2C^+^FcRγ^-/low^ and CD56^bright^CD16^-^ cells (Fig. 3A). Minor variations in CD122 or CD132 levels were detected between CD57^-^ cells and CD57^+^ cells within each mature NK cell subset (Fig. 3B). Therefore, we concluded that FcRγ expression is not regulated directly by CD122 expression or by the CD57^+^ phenotype at steady-state.

**Figure 3:**
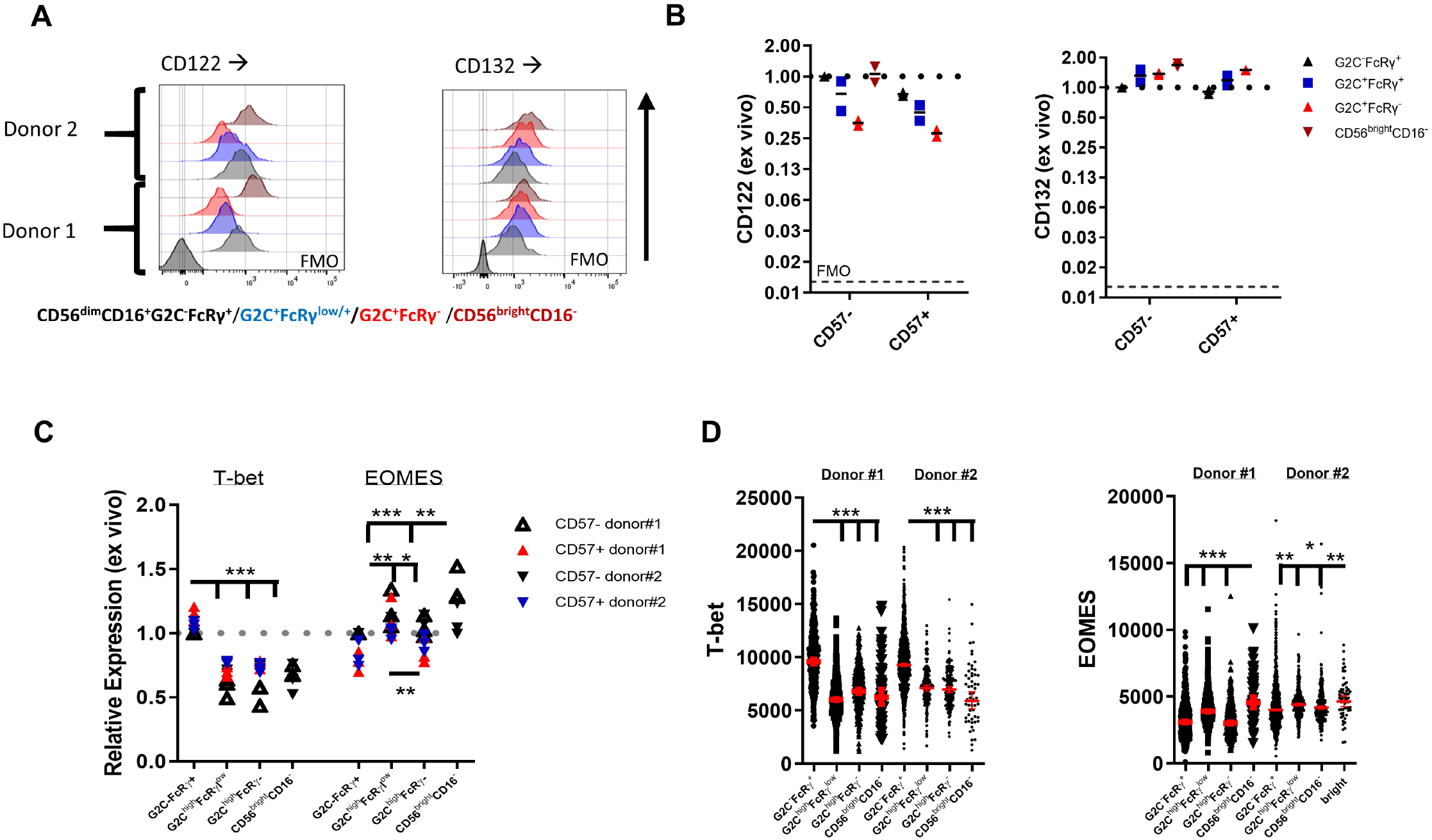
CD122 expression is coupled to FcRγ levels. **(A)** Representative histograms of CD122 (IL-2R) and CD132 (IL-2Ry) expression on CD56^bright^CD 16^-^ and the specified CD56^dim^CD16^+^ NK cell subsets, *ex vivo*. (FMO = fluorescence minus one control). **(B)** Surface CD122 or CDl32 gMFT levels on CD56^bright^CD16^-^ and the specified CD56^dim^CD16^+^ NK cell subsets, *ex vivo*. Values are normalized to CD57^-^NKG2C^-^FcRγ^+^ (index = 1). **(C)** Intracellular T-bet or EOMES levels in the defined NK cell subsets, Values were normalized to CD56^dim^CD16^+^NKG2C^-^FcRγ^+^CD57^-^ NK cells (index = 1). Each measurement represents independent staining of an independent blood sample collected at an independent time point. **(D)** Single cell expression of T-bet or EOMES in the defined NK subsets, linear values. Red bar = geometric mean with 95% confidence interval, unpaired t-test, two tails, *p<0.05, **p<0.01, *** p<0.001.

CD122 expression is suggested to be regulated by EOMES and T-bet in immature and mature NK cells, respectively, and are regulated downstream of mTOR activity ^43^. Therefore, we analyzed EOMES and T-bet levels relative to FcRγ expression. T-bet expression was lower in CD56^dim^NKG2C^+^FcRγ^-^, CD56^dim^NKG2C^+^FcRγ^low^, and CD56^bright^CD16^-^ NK cells (Fig. 3C, 3D). EOMES expression was higher in CD56^bright^CD16^-^ cells and showed variable expression between mature NK cell subsets. Therefore, we concluded FcRγ levels might be regulated similar to T-bet expression.

### Rapamycin or TGFβ inhibit FcRγ upregulation

FcRγ expression in NK cells is reduced by CD16 stimulation and restored by IL-2 stimulation ^19^, yet the mechanism of FcRγ upregulation in NK cells is not well defined. As our observations suggest FcRγ is regulated by mTOR activity, we stimulated purified NK cells from 3 donors with IL-2 in the presence of rapamycin or TGFβ and evaluate FcRγ expression. Either TGFβ or rapamycin significantly inhibited FcRγ upregulation during IL-2 stimulation (Fig 4A, 4B). Cell division progression, measured as reduced cell tracer violet (CTV) signal, significantly correlated with FcRγ upregulation (Fig. 4C, 4D). The increase in FcRγ levels was not associated with a specific NK cell subset, and higher IL-2 dose stimulation reduced the frequencies of the mature NKG2C^high^FcRγ^-^ NK cells (Fig. 4E), which was inhibited by rapamycin even during CD16 stimulation. Interestingly, rapamycin did not influence CD25 upregulation, while NK cell size did not differ between the indicted conditions (Fig S3). Therefore, we concluded that FcRγ protein levels are regulated by mTOR activity and might be coupled to NK cell division progression.

**Figure 4:**
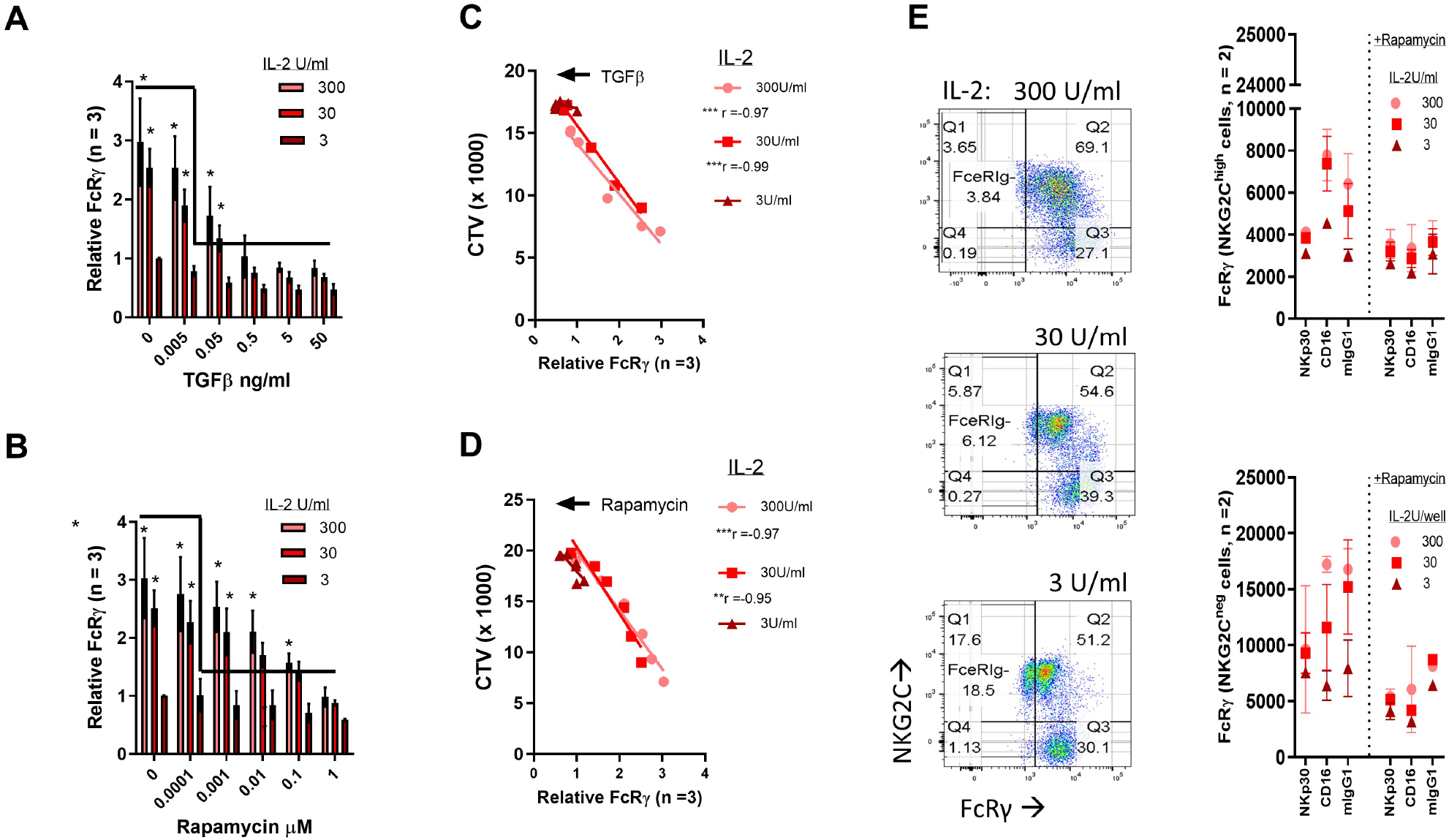
Rapamycin or TGFβ inhibit FcRγ upregulation and cell division. **(A-B)** FcRγ upregulation by IL-2 stimulation in the presence of variable TGFβ **(A)** or rapamycin **(B)** concentrations. Integrated data from three independent experiments (n = 3), n = I/experiment, mean± S.D, paired t-test, two-tails, **p<0.01, ***p<0.001 (nl= donor #1: NKG2C^high^, Donor#3 and donor#4 = donors lacking NKG2C^high^ expression or the CD56^dim^FcRγ^-^ subset. FcRγ expression was nonnalized to FcRγ levels in the presence of IL-2 (3 U/ml) stimulation without TGFB or rapamycin (paired t-test, two-tails, *p<0.05) **(C-D)** Correlation between mean FcRγ gMFI levels and mean CTV gMFI levels (n = 3) in the presence of TGFβ (C) or rapamycin (D), Pearson correlation, two-tails, **p<0.01, ***p<0.001). **(E)** Representative dot plot of FcRγ vs. NKG2C expression on CD56^dim^CD16^+^ cells following IL-2 stimulation (day 6). (Donor #1). Right: FcRγ gMFI levels in NKG2C^high^FcRγ^+/-^ (upper) or NKG2C-FcRγ^+^ (lower) NK cells following the indicted stimulation in the presence or absence of rapamycin (0.1 µM) (day 5). Mean± S.D Integrated results from donors #1 and #2.

### FOXO1 regulates FcRγ expression downstream of mTOR activity

FOXO1 function regulates EOMES and T-bet expression as well as NK cell division downstream of mTOR activity ^35,40,56^. Additional, FOXO1 suppresses cell division progression and is negatively regulated by mTOR activity^33^. To evaluate if FOXO1 activity regulates FcRγ levels, we purified NK cells from 6 donors without CD56^dim^NKG2C^high^ or/and CD56^dim^FcRγ^-/low^ subsets and stimulated them with high IL-2 doses in the presence of rapamycin and a FOXO1 inhibitor (FOXO1i; AS1842856 IC_50_ = 33 nM). Co-culture of primary NK cells with rapamycin and FOXO1i (50 nM) increased cell division and FcRγ expression mediated by IL-2 stimulation (Fig.5A-C). Furthermore, T-bet expression and the percentage of Ki-67^+^ NK cells were suppressed by rapamycin and significantly upregulated by inhibition of FOXO1 (Fig. 5D, 5E). Although both T-bet and FcRγ were upregulated during FOXO1 inhibition, FcRγ upregulation did not significantly correlate to T-bet expression but did show a significantly positive correlation to the percentage of Ki-67^+^ cells during NK cell stimulation (Fig. 5F, 5G). In the presence of TGFβ, FOXO1 inhibition increased NK cell division, FcRγ levels, and the percentage of Ki-67^+^ cells, yet the effects of FOXO1 inhibitor were more variable on TGFβ relative to rapamycin, suggesting a differential TGFβ sensitivity between donors or the involvement of Smad proteins in cell division regulation and FcRγ expression ^33^ (Fig. 5H-J).

**Figure 5:**
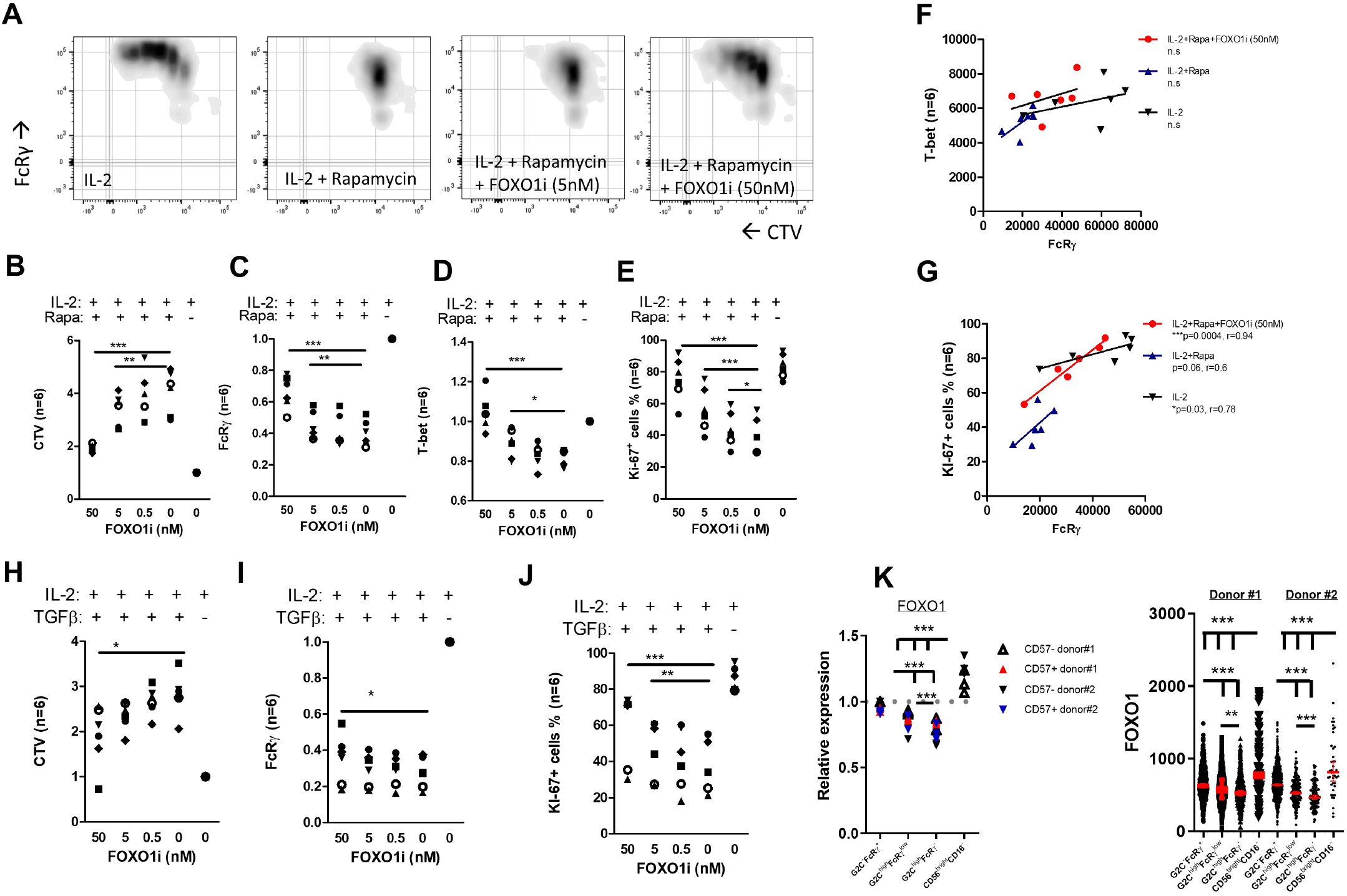
FOXOl inhibition leads to FcRγ upregulation in the presence of rapamycin or TGFβ. **(A)** Representative density plot of FcRγ expression vs. CTV in CD56^+^CD16^-/+^ NK cells following 7 days of IL-2 stimulation (300 U/ml). **(B-E)** CTV **(B)**, FcRγ (C), T-bet **(D)** gMFI levels, or percentage of Ki-67^+^ cells **(E)** in CD56^+^CD16^+^CD16^-/+^ NK cells following 7 days of IL-2 stimulation in the presence or absence of rapamycin (0.1 µM) and the specified FOXOl inhibitor concentrations. Values were normalized to IL-2 stimulation without inhibitors (n= 6, donors# 5-10, each symbol = 1 donor, paired t-test (two-tails, *p<0.05, **p<0.01, ***p<0.001). **(F-G)** Correlation between FcRγ and T-bet expression **(F)** or FcRγ and percentage ofKi-6r cells **(G)** (n= 6, Pearson correlation, one tail). **(H-J)** CTV levels **(H)**, FcRγ levels (I), or percentage of K.i-6r cells **(J)** in CD56^+^CDI6-1^+^ NK cells following 7 days of lL-2 stimulation in the presence or absence of TGF (0.5 ng/ml) and the specified FOXOl inhibitor concentrations. Values were normalized to IL-2 stimulation without inhibitors (n= 6, each symbol = I donor, paired t-test (two-tails, *p<0.05, **p<0.01, ***p<0.001). **(K)** Intracellular FOXOI levels in CD56^bright^CD16^-^ and the specified CD56^dim^CD16^+^ NK cells, *ex vivo*. gMFI values were normalized to NKG2C^-^ FcRγ^+^CD57-cells (index = 1). Each measurement represents independent staining obtained from a blood sample that was collected at an independent time point. Paired t-test, two tails, *p<0.05, **p<0.01, ***p<0.001. Right: Single cell expression of FOXOl in the defined NK subsets, linear values. Red bar = geometric mean with 95% confidence interval, unpaired t-test, two tails, **p<0.0 1, *** p<0.001.

To evaluate if FOXO1 levels are associated with FcRγ levels *ex vivo* at steady-state, we examined FOXO1 levels in NK cell subsets from donor #1 and #2 (Fig. 2). At steady-state FOXO1 levels were significantly lower in CD56^dim^NKG2C^high^FcRγ^-^ or FcRγ^low^ NK cells, while immature CD56^bright^CD16^*-*^ cells expressed higher levels (Fig. 5K). Therefore, we concluded that FcRγ levels are regulated by NK cell division progression downstream of mTOR activity and FOXO1 function.

## Discussion

Viral infections can lead to the accumulation of CD56^dim^CD16^+^NKG2A^+/-^NKG2C^+/-^FcRγ^-/low^ NK cells, yet the molecular mechanisms regulating FcRγ levels are not well defined. As this adaptor protein regulates the function of NKp30, NKp46, and CD16, which are shown to take part in anti-viral responses, cancer immunosurveillance, and tumor rejection, it is important to better characterize the mechanism of FcRγ regulation^8^.

Here we have discovered that FcRγ expression corresponds to cell division progression in NK cells downstream of mTOR activity. Therefore, FcRγ levels *in vivo* might reflect NK cell division suppression. As CD16 was reported to mediate FcRγ loss during NK cell activation, we suggest here that NK cell division progression following CD16 stimulation (or other activating receptors), is required to upregulate and sustain FcRγ protein levels. This conclusion is supported by our observations in: 1. lung transplant patients treated with rapamycin, 2. the association of FcRγ levels with mTOR activity in adaptive FcRγ^-/low^ and immature NK cells in healthy donors, 3. the inhibition of FcRγ upregulation during IL-2 stimulation by rapamycin or TGFβ, and 4. by the upregulation of FcRγ via inhibiting FOXO1 activity during IL-2 and rapamycin or TGFβ co-culture.

Although we have shown here that mTOR activity is important for NK cell division and FcRγ upregulation, the IL-2/IL-15 axis might also regulate FcRγ levels by STAT5, which is functionally inhibited by FOXO1and regulate NK cell diviosn^33^; however, our results indicate cell division as a significant regulator of FcRγ expression in NK cells. This also suggests that NK cell division by IL-12 + IL-18, which was documented during mouse cytomegalovirus infection, would lead to FcRγ upregulation in human NK cells similar to observations seen for mouse PLZF^50^. Additionally, lower Bcl2 levels correlate with cell cycle progression in mouse NK cells, which might suggest human adaptive FcRγ^-/low^ NK cell persist by reducing NK cell division associated with lower mTOR activity yet without affecting STAT5 signaling^21^. A second possibility is a slow proliferation maintained by receptor stimulation of adaptive FcRγ^-/low^ NK cells would sustain low FcRγ levels, yet we did not find evidence supporting this hypothesis. Additionally, observations in T cells, which are usually negative for FcRγ, show *FCER1G* promotor demethylation by IL-15 whereas the transcription factor SP1, a positive regulator of the *FCER1G* promotor, is regulated by cell cycle-related proteins^57–59^. Additional possibility is regulation of mTOR activity by FcRγ expression, as was reported in mast cells, yet as NK cells express additionally adaptor proteins such as CD3ζ and DAP12, this might compensate the loss of FcRγ^13^.

FcRγ levels were associated with T-bet expression in healthy donors, and both were upregulated during cell division progression. FOXO1-SP1 interaction regulates T-bet expression in NK cells^56^. T-bet is associated with short-live effector CD8^+^ T cells while being suppressed to promotes memory CD8^+^ T cells differentiation and maintenance during chronic viral infections to prevent anergy ^35,60,61^. This indicates that lower FcRγ levels may define memory or exhaustion in human NK cells. In our analysis, immature CD56^bright^CD16^-^ NK cells show a signaling profile resembling memory CD8^+^ T cells, with FOXO1^high^/EOMES^high^ and mTOR^low^/T-bet^low^ expression but maintain their ability to proliferate by high CD122 expression important to activate mTOR^39^. However, adaptive CD56^dim^FcRγ^-/low^ NK cells were FOXO1^low^EOMES^+^ and mTOR^low^/T-bet^low^/Bcl2^high^; therefore, showing a distinct NK cell “memory” signaling differentiation resembling effector-memory or exhausted CD8^+^ T cells ^60,62^. In T cells, higher FcRγ levels are reported to associated with higher anti-tumor activity^57^. As we report here, CD16 stimulation led to upregulation of FcRγ in a IL-2 dependent manner, yet additional work is required to examine if this exhaustion/memory signature of adaptive CD56^dim^FcRγ^-/low^ NK cells during HCMV reactivation is associated with NK cell division and FcRγ upregulation.

The interest in human NK cells and more specifically human adaptive CD56^dim^FcRγ^-/low^ NK cells in cancer immunotherapy is rising as these cells do not express the inhibitory receptor NKG2A and have greater persistence ^63,64^. The tumor microenvironment is rich with NK cell immunosuppressive factors such as TGFβ ^36,37^. We have shown here that TGFβ, like rapamycin, prevented FcRγ upregulation during high-dose IL-2 stimulation, which is significantly correlated with NK cell division progression. As FcRγ regulates the expression of NKp30, NKp46 and CD16, this mechanism of FcRγ suppression might promote tolerance as stimulation of activating receptor can enhance mTOR activation by pro-inflammatory cytokines^5,11,65^. In acute myeloid leukemia (AML), the loss of NKp30 and NKp46 on NK cells is associated with a poor prognosis ^9,10,20^. Additionally, reduced NKp30 or NKp46 levels on NK cells are reported during pregnancy, which is associated with maternal-fetal immune tolerance^66^. Additionally, low mTOR activity in human and mouse NK cells is reported to diminish NK cell effector functions against tumors *in vivo* ^67^. As adaptive CD56^dim^FcRγ^-/low^ NK cell frequencies are stable over time, this might suggest the need to develop approaches to target adaptive FcRγ^-/low^ NK cells to increase NK cancer immunosurveillance or to engineer these cells for higher pro-inflammatory cytokine sensitivity to enhance their intratumoral functions.

Our findings present new insights on FcRγ regulation in human NK cells and might contribute to our understating on the differentiation and development of adaptive CD56^dim^FcRγ^-/low^ NK cells, and to the development of new therapeutic approaches to enhance NK cell functions in cancer immunosurveillance and cancer immunotherapy.

## Materials and Methods

### Lung transplant recipients samples

The UCSF Institutional Review Board (IRB# 13-10738) approved this study. Lung transplant recipient standard maintenance immunosuppressant therapy included tacrolimus, prednisone, and mycophenolate mofetil. Tacrolimus troughs of 8-14 ng/ml were targeted during the first 12 months after transplant and 6-10 ng/ml thereafter. All subjects were started on 20 mg of prednisone daily and reduced over the first post-operative year to a goal dose of 0.1 mg/kg. Sirolimus is used as adjunctive therapy as a treatment for chronic lung allograft dysfunction (CLAD) or recurrent episodes of acute cellular rejection (ACR). In addition, sirolimus is prescribed as a calcineurin-inhibitor sparing agent when subjects developed chronic kidney disease (CKD) or recurrent skin carcinomas. PBMC were prospectively collected and cryopreserved from lung transplant recipients during routine clinic visits. Four subjects were identified with PBMC available before and during therapeutic sirolimus treatment, which was targeted through blood drug levels. PBMCs were identified in five control lung transplant subjects that were at least 1-year post-transplant and matched for transplant indication.

### Primary NK cells isolation and culture

Primary human NK cells were obtained from healthy donors’ peripheral blood after donors gave informed consent in accordance with approval by the UCSF Institutional Review Board (IRB# 10 - 00265) or from Plateletpheresis leukoreduction filters (Vitalant, https://vitalant.org/Home.aspx). NK cell isolation was done by using the negative selection kit “RosetteSep™ Human NK Cell Enrichment Cocktail” (STEMCELL Technologies) according to the company’s protocol. Purified NK cells (CD56^+^CD3^-^) were used on the same day (day 0, *ex vivo*) or after priming with IL-2 as indicated. NK cells culture media: GMP SCGM (CellGenix®) supplemented with 1% L - glutamine, 1% penicillin and streptomycin, 1% sodium pyruvate, 1% non-essential amino acids, 10 mM HEPES, and 10% human serum (heat-inactivated, sterile-filtered, male AB plasma; SIGMA). Human IL-2 (TECIN™ teceleukin, ROCHE, was generously provided by the NCI Biological Resources Branch. Donor #1 HLA-C genotype (Cw*04 and Cw*06).

### Antibody-conjugated beads antibody and biotinylation cytokines, and inhibitors

Antibody-conjugated beads were prepared according to the company’s protocol (Invitrogen™ Dynabeads™ Antibody Coupling Kit) at 10 µg antibody for 1 mg beads. Following conjugation, beads were resuspended in sterile PBS at an antibody concentration of 0.1 µg/µl. Antibody conjugation was evaluated by flow cytometry with APC-conjugated anti-mouse or -rat IgG. BioLegend: anti-human CD16 (cat. 302002, mouse IgG1k), anti-human NKp30 (cat. 325204, mouse IgG1k). UCSF Monoclonal Antibody Core: mouse IgG1 isotype matched-control (clone; MOPC-21). Antibody biotinylation was preform using an EZ-Link Micro NHS-PEG4 Biotinylation kit (ThermoFisher cat. 21955).

### Cytokines and inhibitors

All cytokines were resuspended in sterile PBS: human IL-2, 1,000 U/µl (TECINTM; teceleukin, ROCHE, generously provided by NCI Biological Resources Branch); human TGFβ1, 50 µg/ml (BioLegend, cat. 580706). All inhibitors were resuspended in DMSO; mTORC1 inhibitor (Calbiochem; Rapamycin, cat. 553210, IC_50_ = 0.1 µM), FOXO1 inhibitor Calbiochem; (AS1842856, cat. 344355, IC_50_ = 33 nM). Antibody-conjugated beads were stored at 4°C. Cytokines or inhibitors were stored at-20° C.

### NK cell stimulation and cell division assay

*Ex vivo* NK cells were labeled with cell tracer violet dye (CTV) according to t he company’s protocol (Invitrogen™, cat. C34557). The final culture volume during the assay was 200 µl/well in 96-well plate round-bottom plates. Antibody-coated beads were diluted 1:500 from stock (0.1 µg/µl) in cytokine-free NK cell culture media to an antibody concentration of 0. 2 ng/µl and 50 µl were used during the assay. The final antibody amount per well during the assay was 0.01 µg per well. IL-2 was diluted to a final concentration of 1200, 120, or 12 U/ml and 50 µl were used during the assay. Final IL-2 concertation during the assay was 300, 30, or 3 U/ml. The final IL-2 amount per well was 60, 6, or 0.6 U/well. TGFβ1 was diluted to 200, 20, 2, 0.2, 0.02 ng/ml and 50 µl were used during the assay at a final concentration of 50, 5, 0.5, 0.05, 0.005 ng/ml. Final TGFβ1 amount per well was 10, 1, 0.1, 0.01, 0.001 ng/well. Rapamycin (Calbiochem; Rapamycin, cat. 553210) or AS1842856 (FOXO1i) were used at the indicated final concentrations. Cytokine-free NK cell culture media was added at 50-100 µl/well to reach a culture volume of 150 µl/well. At the final step, CTV-labeled NK cells were added at 5 × 10^4^ cells/well, 50 µl/well, and incubated at 37°C with 5% CO_2_ for the duration of the assay. CTV signal was analyzed by flow cytometry (LSR-II, Becton Dickinson Immunocytometry Systems). For cell division assays without activating receptor stimulation, CTV signal was analyzed at day 6, with activating receptor stimulation at day 5.

#### CD25, CD107a, and pS6 assays

NK cells (5 × 10^4^ cells/well) were stimulated for 24 hours with antibody-coated beads (0.01 µg per well) + IL-2(300-30-3 U/ml) at a final volume of 200 µl with or without rapamycin (1µM) as indicated. AF-647-conjugated anti-CD107a (BioLegend, cat. 328612) was added at the beginning of the assay to the cell culture at a final dilution of 1:100 and by the end of the assay during membrane staining (1:200, 50 µl/well, 30 min, 4 °C). In some cases, FcRγ levels were normalized to minimized expression levels differences between donors and between integrated experiments.

### Flow cytometry

*Ex vivo* NK cells (5 × 10^4^ cells/well) and NK cells following specified stimulation and at the indicated time points were collected and culture media was washed out using flow buffer (PBS +2% FCS). For membrane staining, antibodies were resuspended in PBS + 2% FCS and incubated with the cells for 30 min at 4 °C at 50 µl/well. Dead cells were excluded by labeling using PI (1 mg/ml, 1:500) or near IR-fixable dye (1:1000, Invitrogen, cat. L34976). For the wash step, 150 µl/well of flow buffer was added, the plate was centrifuged at 600 x g rcf for 5 min at 4 °C, and the buffer was discarded. For intracellular expression, cells were incubated for 20 min at 4 °C with 100 µl/well Cytofix/Cytoperm buffer (Becton Dickinson, cat. 51-2090KZ). Following incubation, cells were washed twice using 150 µl/well intracellular-staining Perm-Wash buffer (BioLegend, cat. 421002) diluted 1:10 in PBS. Antibodies against intracellular markers were diluted in Perm-Wash buffer and incubated with the cells for 60 min at 4 °C. Following incubation, cells were washed twice with 150 µl/well Perm-Wash buffer. Before analyzing the samples for membrane or/and intracellular markers, cells were resuspended in a 300 µl flow buffer. Samples were kept at 4°C until analyzed using an LSR-II flow cytometer.

### Antibodies against membrane or intracellular proteins

For detection of NK cell subsets, as indicated, the following antibodies were used: BioLegend: APC-Cy7-conjugated anti-CD3 (Cat# 300318, RRID:AB_314054), PerCep-Cy5.5 anti-CD56 (Cat# 318322, RRID:AB_893389), APC-conjugated anti-CD16 (Cat# 302012, RRID:AB_314212), or BV-421-conjugated anti-CD16 (Cat# 302038, RRID:AB_2561578) or PE-Cy7-conjugated anti-CD16 (Cat# 360708, RRID:AB_2562951), BV605-conjugated anti-CD57 (Cat# 393304, RRID:AB_2728426) or APC-conjugated anti-CD57 (Cat# 322314, RRID:AB_2063199), R&D Systems, PE- or APC-conjugated anti-NKG2C (Cat# FAB138P, RRID:AB_2132983 or Cat# FAB138A, RRID:AB_416838 – generously provided by R&D Systems), and then stained for intracellular expression with FITC anti-FcRγ (Millipore Cat# FCABS400F, RRID:AB_11203492). For detection of surface or intracellular proteins: <u>BioLegend</u>: APC-conjugated anti-NKp30 (Cat# 325210, RRID:AB_2149449), APC-conjugated anti-NKp46 (Cat# 331918, RRID:AB_2561650), APC-conjugated anti-CD25 (at# 302610, RRID:AB_314280), APC-conjugated anti-CD122 (Cat# 339008, RRID:AB_2123575), PE-conjugated anti-CD132 (Cat# 338605, RRID:AB_1279079), APC-conjugated anti-DNAM1 (Cat# 338312, RRID:AB_2561952). APC-conjugated anti-human IL-2 (Cat# 500310, RRID:AB_315097), AF-647 -conjugated anti-IFNγ (Cat# 502516, RRID:AB_493031), and APC--conjugated anti-SYK (Cat# 644305, RRID:AB_2687145), APC-conjugated anti-T-bet (Cat# 644813, RRID:AB_10896913), AF647-conjugated anti-CD107b (Cat# 354311, RRID:AB_2721404), APC-conjugated anti-ADAM10 (Cat# 352705, RRID:AB_2563172) APC-conjugated streptavidin (cat. 405207). <u>R&D Systems</u>: PE-conjugated anti-KIR2DL1 (Cat# FAB1844P, RRID:AB_2130401), PE-conjugated anti-KIRD2L2/3 (clone DX27), PE-conjugated anti-KIR3DL1 (clone DX9), and PE-conjugated anti-KIR3DL2 (clone DX31), AF647-conjugated anti-LAT1 (cat. FAB10390R), AF647-conjugated anti-Glut1 (cat. FAB1418R), APC-conjugated anti-CD98 (Cat# FAB5920A, RRID:AB_1964536), AF647-conjugated anti-PLZF (cat. IC2944R) and APC-conjugated anti-IL-15 (Cat# IC2471A, RRID:AB_10714827), AF647-conjugated anti EOMES (cat. IC6166R), AF647-conjugated anti-RPTOR (cat. IC5957R), purified sheep-anti-Rictor (at# AF4598, RRID:AB_2179841).<u>IOTest</u>: PE-conjugated anti-human CD85J (cat. PN A07408). <u>Becton Dickinson:</u> APC-conjugated anti-CD122 (Mik-β3, cat. Cat# 566620, RRID:AB_2869796), AF647-conjugated anti-pSTAT4-Y693 (Cat# 562074, RRID:AB_10896660), AF647-conjugated anti-pSTAT5-Y694 (Cat# 612599, RRID:AB_399882), AF647-conjugated anti-pSTAT1-Y701 (Cat# 612597, RRID:AB_399880), AF647-conjugated anti-Bcl-2 (Cat# 563600, RRID:AB_2738306), AF647-conjugated anti-Ki-67 (Cat# 558615, RRID:AB_647130). <u>Cell Signaling</u>: AF647-conjugated anti-pS6-S235/236 (Cat# 4851, RRID:AB_10695457), AF647-conjugated anti-pAKT-T308 (Cat# 48646, RRID:AB_2799341), AF647-conjugated anti-pAKT-S473 (Cat# 4075, RRID:AB_916029), AF647-conjugated anti-mTOR (at# 5048, RRID:AB_10828101), AF647-conjugated anti FOXO1 (Cat# 72874, RRID:AB_2799829), purified anti-pRptor S792 (cat. 89146S, biotinylated), AF647 anti pp38-T180/Y182 (Cat# 4552, RRID:AB_331303)<u>.Miltenyi-Biotech:</u> APC-conjugated anti-NKG2A (Cat# 130-114-089, RRID:AB_2726447), PE-vio-770-conjugated anti NKG2A (cat. Cat# 130-113-567, RRID:AB_2726172). Invitrogen: AF647-conjugated donkey anti-sheep IgG (at# A21448, RRID:AB_1500712).

### Flow-cytometry analysis of lung transplant patients

Lymphocytes in the PBMC were gated as CD45^+^ (BioLegend cat. 304024), live cells (Zombie red-negative, BioLegend cat.77475), and singlet cells based on forward-angle-light-scattering properties (FSC-A vs. FSC-H). Non-NK cells were excluded by staining with anti-CD3 (BioLegend Cat# 300318, RRID:AB_314054), -CD19 (BioLegend Cat# 302218, RRID:AB_314248), -CD4 (BioLegend Cat# 357416, RRID:AB_2616810), -CD14 (BioLegend Cat# 301820, RRID:AB_493695), and -CD123 (BioLegend Cat# 306042, RRID:AB_2750163). NKp46^+^ cells were gated as NKG2A vs. CD16. NKG2A-negative CD16-negative cells were excluded as they could not be defined as a specific NK cell subset. Antibodies used included anti-NKp46 (BioLegend Cat# 331936, RRID:AB_2650940), -CD16 (BioLegend Cat# 302038, RRID:AB_2561578), and PE-vio-770-conjugated anti-NKG2A (cat. Cat# 130-113-567, RRID:AB_2726172). Human Fc receptors were blocked using human TruStain FcXTM (BioLegend Cat# 422302, RRID:AB_2818986).

### Graphics and statistical analysis

Graphs were generated using GraphPad Prism 5 or FlowJo_V10. Statistical analysis as indicated in figure legends was calculated using GraphPad Prism 5 or Excel 2013. p*<0.05, p**<0.01, p***<0.001. Statistical tests are described in figure legends.

## Author Contributions

Conceptualization, A.S. and L.L.L.; Methodology, A.S.; Investigation, A.S. and J.H.A.; Formal Analysis, A.S.; Resources, D.R.C., and J.R.G.; Funding Acquisition, L.L.L, A.S., D.D.C., J.R.G.; Writing – Original Draft, A.S. and L.L.L.; Writing – Review & Editing, D.R.C., and J.R.G.; Supervision, L.L.L. The authors declare no competing interests.

## Acknowledgments

Studies were supported by NIH grant AI068129, the Parker Institute for Cancer Immunotherapy, and the Irvington Cancer Research Institute Fellowship to A.S, the Joel D. Cooper Award from the International Society for Heart and Lung Transplantation (D.R.C), the Cystic Fibrosis Foundation Harry Shwachman Career Development Award CALABR19Q0 (D.R.C), the Veterans Affairs Office of Research and Development (CX002011, J.R.G) and NHLBI (HL151552, J.R.G). We acknowledge Emily Aminian, Lily Tran, Jon Singer, Steve Hays, and the remainder of the lung transplant clinical team for instrumental aid in subject recruitment and sample processing. We are thankful to the organ and tissue donors, and their families for giving gifts of life and knowledge with their generous donation, the UCSF Parnassus Flow Core (RRID: SCR_018206 and DRC Center Grant NIH P30 DK063720).

## References

1. Cerwenka, A. & Lanier, L. L. Natural killer cell memory in infection, inflammation and cancer. Nat Rev Immunol 16, 112–123 (2016).

2. Cooper, M. A., Fehniger, T. A. & Caligiuri, M. A. The biology of human natural killer-cell subsets. Trends in Immunology 22, 633–640 (2001).

3. Björkström, N. K. et al. Expression patterns of NKG2A, KIR, and CD57 define a process of CD56dim NK-cell differentiation uncoupled from NK-cell education. Blood 116, 3853–3864 (2010).

4. Lopez-Vergès, S. et al. CD57 defines a functionally distinct population of mature NK cells in the human CD56dimCD16+ NK-cell subset. Blood 116, 3865–3874 (2010).

5. Lanier, L. L., Yu, G. & Phillips, J. H. Analysis of Fc gamma RIII (CD16) membrane expression and association with CD3 zeta and Fc epsilon RI-gamma by site-directed mutation. J Immunol 146, 1571–1576 (1991).

6. Muccio, L. et al. Late Development of FcεRγneg Adaptive Natural Killer Cells Upon Human Cytomegalovirus Reactivation in Umbilical Cord Blood Transplantation Recipients. Front. Immunol. 9, (2018).

7. Hart, G. T. et al. Adaptive NK cells in people exposed to Plasmodium falciparum correlate with protection from malaria. Journal of Experimental Medicine 216, 1280–1290 (2019).

8. Sheffer, M. et al. Genome-scale screens identify factors regulating tumor cell responses to natural killer cells. Nat Genet 1–11 (2021) doi:10.1038/s41588-021-00889-w.

9. Chretien, A.-S. et al. NKp46 expression on NK cells as a prognostic and predictive biomarker for response to allo-SCT in patients with AML. null 6, e1307491 (2017).

10. Chretien, A.-S. et al. NKp30 expression is a prognostic immune biomarker for stratification of patients with intermediate-risk acute myeloid leukemia. Oncotarget 8, 49548–49563 (2017).

11. Liu, W. et al. FcRγ Gene Editing Reprograms Conventional NK Cells to Display Key Features of Adaptive Human NK Cells. iScience 23, 101709 (2020).

12. Lee, J. et al. Epigenetic Modification and Antibody-Dependent Expansion of Memory-like NK Cells in Human Cytomegalovirus-Infected Individuals. Immunity 42, 431–442 (2015).

13. Saitoh, S. et al. LAT is essential for Fc(epsilon)RI-mediated mast cell activation. Immunity 12, 525–535 (2000).

14. Hammer, Q. et al. Peptide-specific recognition of human cytomegalovirus strains controls adaptive natural killer cells. Nature Immunology 19, 453–463 (2018).

15. Lopez-Vergès, S. et al. Expansion of a unique CD57+NKG2Chi natural killer cell subset during acute human cytomegalovirus infection. PNAS 108, 14725–14732 (2011).

16. Liu, L. L. et al. Critical Role of CD2 Co-stimulation in Adaptive Natural Killer Cell Responses Revealed in NKG2C-Deficient Humans. Cell Reports 15, 1088–1099 (2016).

17. Gyurova, I. E. et al. Dynamic Changes in Natural Killer Cell Subset Frequencies in the Absence of Cytomegalovirus Infection. Front. Immunol. 10, (2019).

18. Schlums, H. et al. Cytomegalovirus Infection Drives Adaptive Epigenetic Diversification of NK Cells with Altered Signaling and Effector Function. Immunity 42, 443–456 (2015).

19. Pahl, J. H. W. et al. CD16A Activation of NK Cells Promotes NK Cell Proliferation and Memory-Like Cytotoxicity against Cancer Cells. Cancer Immunol Res 6, 517–527 (2018).

20. Martner, A. et al. NK cell expression of natural cytotoxicity receptors may determine relapse risk in older AML patients undergoing immunotherapy for remission maintenance. Oncotarget 6, 42569–42574 (2015).

21. Marçais, A. et al. The metabolic checkpoint kinase mTOR is essential for IL-15 signaling during the development and activation of NK cells. Nature Immunology 15, 749–757 (2014).

22. Gasteiger, G. et al. IL-2–dependent tuning of NK cell sensitivity for target cells is controlled by regulatory T cells. J Exp Med 210, 1167–1178 (2013).

23. Anton, O. M., Vielkind, S., Peterson, M. E., Tagaya, Y. & Long, E. O. NK Cell Proliferation Induced by IL-15 Transpresentation Is Negatively Regulated by Inhibitory Receptors. J Immunol 195, 4810–4821 (2015).

24. Anton, O. M. et al. Trans-endocytosis of intact IL-15Rα–IL-15 complex from presenting cells into NK cells favors signaling for proliferation. PNAS 117, 522–531 (2020).

25. Vámosi, G. et al. IL-2 and IL-15 receptor α-subunits are coexpressed in a supramolecular receptor cluster in lipid rafts of T cells. Proc Natl Acad Sci U S A 101, 11082–11087 (2004).

26. Wiedemann, G. M. et al. Divergent Role for STAT5 in the Adaptive Responses of Natural Killer Cells. Cell Reports 33, 108498 (2020).

27. Eckelhart, E. et al. A novel Ncr1-Cre mouse reveals the essential role of STAT5 for NK-cell survival and development. Blood 117, 1565–1573 (2011).

28. Sun, J. C. et al. Proinflammatory cytokine signaling required for the generation of natural killer cell memory. J Exp Med 209, 947–954 (2012).

29. Wang, K. S., Ritz, J. & Frank, D. A. IL-2 Induces STAT4 Activation in Primary NK Cells and NK Cell Lines, But Not in T Cells. The Journal of Immunology 162, 299–304 (1999).

30. Ali, A. K., Nandagopal, N. & Lee, S.-H. IL-15–PI3K–AKT–mTOR: A Critical Pathway in the Life Journey of Natural Killer Cells. Front. Immunol. 6, (2015).

31. Keating, R. & McGargill, M. A. mTOR Regulation of Lymphoid Cells in Immunity to Pathogens. Front. Immunol. 7, (2016).

32. Hart, J. R. & Vogt, P. K. Phosphorylation of AKT: a Mutational Analysis. Oncotarget 2, 467–476 (2011).

33. Calnan, D. R. & Brunet, A. The FoxO code. Oncogene 27, 2276–2288 (2008).

34. Sarbassov, D. D. et al. Prolonged Rapamycin Treatment Inhibits mTORC2 Assembly and Akt/PKB. Molecular Cell 22, 159–168 (2006).

35. Rao, R. R., Li, Q., Bupp, M. R. G. & Shrikant, P. A. Transcription Factor Foxo1 Represses T-bet-Mediated Effector Functions and Promotes Memory CD8+ T Cell Differentiation. Immunity 36, 374–387 (2012).

36. Viel, S., Besson, L., Marotel, M., Walzer, T. & Marçais, A. Regulation of mTOR, Metabolic Fitness, and Effector Functions by Cytokines in Natural Killer Cells. Cancers (Basel) 9, (2017).

37. Viel, S. et al. TGF-β inhibits the activation and functions of NK cells by repressing the mTOR pathway. Sci. Signal. 9, ra19–ra19 (2016).

38. Yang, C. et al. mTORC1 and mTORC2 differentially promote natural killer cell development. eLife 7, e35619 (2018).

39. Wang, F. et al. Crosstalks between mTORC1 and mTORC2 variagate cytokine signaling to control NK maturation and effector function. Nature Communications 9, 4874 (2018).

40. Zhang, L. et al. Mammalian Target of Rapamycin Complex 2 Controls CD8 T Cell Memory Differentiation in a Foxo1-Dependent Manner. Cell Rep 14, 1206–1217 (2016).

41. Pradier, A. et al. Modulation of T-bet and Eomes during Maturation of Peripheral Blood NK Cells Does Not Depend on Licensing/Educating KIR. Front. Immunol. 7, (2016).

42. Gordon, S. M. et al. The Transcription Factors T-bet and Eomes Control Key Checkpoints of Natural Killer Cell Maturation. Immunity 36, 55–67 (2012).

43. Zhang, J. et al. T-bet and Eomes govern differentiation and function of mouse and human NK cells and ILC1. European Journal of Immunology 48, 738–750 (2018).

44. Benichou, G., Yamada, Y., Aoyama, A. & Madsen, J. C. Natural killer cells in rejection and tolerance of solid organ allografts. Current Opinion in Organ Transplantation 16, 47–53 (2011).

45. Calabrese, D. R. et al. Natural killer cells activated through NKG2D mediate lung ischemia-reperfusion injury. J Clin Invest 131, (2021).

46. Koenig, A. et al. Missing self triggers NK cell-mediated chronic vascular rejection of solid organ transplants. Nature Communications 10, 5350 (2019).

47. Augustine, J. J., Bodziak, K. A. & Hricik, D. E. Use of sirolimus in solid organ transplantation. Drugs 67, 369–391 (2007).

48. Calabrese, D. R., Lanier, L. L. & Greenland, J. R. Natural killer cells in lung transplantation. Thorax 74, 397–404 (2019).

49. Béziat, V. et al. NK cell responses to cytomegalovirus infection lead to stable imprints in the human KIR repertoire and involve activating KIRs. Blood 121, 2678–2688 (2013).

50. Beaulieu, A. M., Zawislak, C. L., Nakayama, T. & Sun, J. C. The transcription factor Zbtb32 controls the proliferative burst of virus-specific natural killer cells responding to infection. Nature Immunology 15, 546–553 (2014).

51. Gwinn, D. M. et al. AMPK Phosphorylation of Raptor Mediates a Metabolic Checkpoint. Molecular Cell 30, 214–226 (2008).

52. Jia, R. & Bonifacino, J. S. Lysosome Positioning Influences mTORC2 and AKT Signaling. Molecular Cell 75, 26-38.e3 (2019).

53. Myers, D. R., Norlin, E., Vercoulen, Y. & Roose, J. P. Active Tonic mTORC1 Signals Shape Baseline Translation in Naive T Cells. Cell Rep 27, 1858-1874.e6 (2019).

54. Shi, H. et al. Amino Acids License Kinase mTORC1 Activity and Treg Cell Function via Small G Proteins Rag and Rheb. Immunity 51, 1012-1027.e7 (2019).

55. Jensen, H., Potempa, M., Gotthardt, D. & Lanier, L. L. IL-2-induced expression of the amino acid transporters SLC1A5 and CD98 is a prerequisite for NKG2D-mediated activation of human NK cells. J Immunol 199, 1967–1972 (2017).

56. Deng, Y. et al. Transcription factor Foxo1 is a negative regulator of natural killer cell maturation and function. Immunity 42, 457–470 (2015).

57. Correia, M. P. et al. Distinct human circulating NKp30+FcεRIγ+CD8+ T cell population exhibiting high natural killer-like antitumor potential. PNAS 115, E5980–E5989 (2018).

58. Tapias, A., Ciudad, C. J., Roninson, I. B. & Noé, V. Regulation of Sp1 by cell cycle related proteins. Cell Cycle 7, 2856–2867 (2008).

59. Takahashi, K. et al. Cooperative Regulation of Fc Receptor γ-Chain Gene Expression by Multiple Transcription Factors, Including Sp1, GABP, and Elf-1. J Biol Chem 283, 15134– 15141 (2008).

60. Delpoux, A., Lai, C.-Y., Hedrick, S. M. & Doedens, A. L. FOXO1 opposition of CD8+ T cell effector programming confers early memory properties and phenotypic diversity. PNAS 114, E8865–E8874 (2017).

61. Delpoux, A. et al. Continuous activity of Foxo1 is required to prevent anergy and maintain the memory state of CD8+ T cells. J Exp Med 215, 575–594 (2018).

62. Staron, M. M. et al. The transcription factor FoxO1 sustains expression of the inhibitory receptor PD-1 and survival of antiviral CD8(+) T cells during chronic infection. Immunity 41, 802–814 (2014).

63. Liu, L. L. et al. Harnessing adaptive natural killer cells in cancer immunotherapy. Mol Oncol 9, 1904–1917 (2015).

64. Creelan, B. C. & Antonia, S. J. The NKG2A immune checkpoint - a new direction in cancer immunotherapy. Nat Rev Clin Oncol 16, 277–278 (2019).

65. Marçais, A. et al. High mTOR activity is a hallmark of reactive natural killer cells and amplifies early signaling through activating receptors. eLife 6, e26423 (2017).

66. Comins-Boo, A. et al. Functional NK surrogate biomarkers for inflammatory recurrent pregnancy loss and recurrent implantation failure. American Journal of Reproductive Immunology n/a, e13426 (2021).

67. Michelet, X. et al. Metabolic reprogramming of natural killer cells in obesity limits antitumor responses. Nature Immunology 19, 1330–1340 (2018).

