## Supplemental figures 1-3 for "Natural killer cell division regulates FcεRIγ expression downstream of mTOR activity"

### Figure S1

**A**

Patients' information table:

| Subject | Transplant indication | Sex | Age | Medications at baseline | Sirolimus indication | Medications post-sirolimus | Sirolimus trough (ug/L) |
|---|---|---|---|---|---|---|---|
| Sirolimus A | ILD | F | 57 | T, MMF, P | CKD | T, P, sirolimus | 6.7 |
| Sirolimus B | ILD | F | 55 | T, MMF, P | CLAD | T, P, sirolimus | 11.1 |
| Sirolimus C | ILD | M | 50 | T, MMF, P | ACR | T, P, sirolimus | 6.3 |
| Sirolimus D | ILD | M | 69 | T, MMF, P | CKD | T, P, sirolimus | 7.1 |
| Control A | ILD | M | 45 | T, MMF, P | | | |
| Control B | ILD | M | 59 | T, MMF, P | | | |
| Control C | ILD | M | 64 | T, MMF, P | | | |
| Control D | ILD | M | 40 | T, MMF, P | | | |
| Control E | ILD | F | 58 | T, MMF, P | | | |

Abbreviations: ILD, interstitial lung disease; M, male; F, female; T, tacrolimus; MMF, mycophenolate mofetil; P, prednisone; CKD, chronic kidney disease; CLAD, chronic lung allograft dysfunction; ACR, acute cellular rejection

**B**

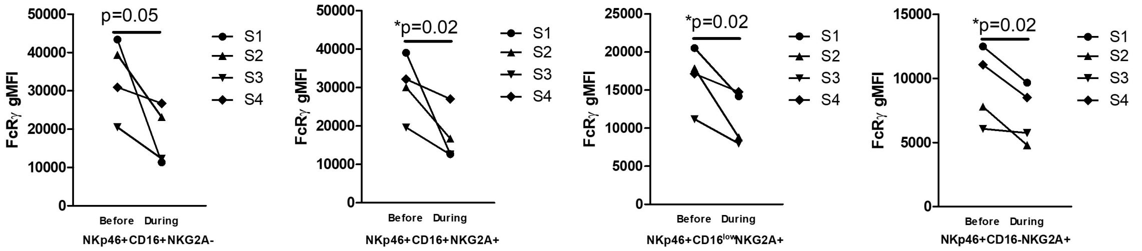


**Supplementary figure 1: Lung transplant recipient clinical information**

**(A)** Table of lung transplant patients' clinical status.

**(B)** FcRγ levels in specified NK cell subsets before or during rapamycin treatment per patient. NK cells were gated as $NKp46^{+}CD3^{-}$ $CD19^{-}CD14^{-}CD4^{-}CD123^{-}$ $CD45^{+}$ live cells. Paired t-test, one-tail.

### Figure S2

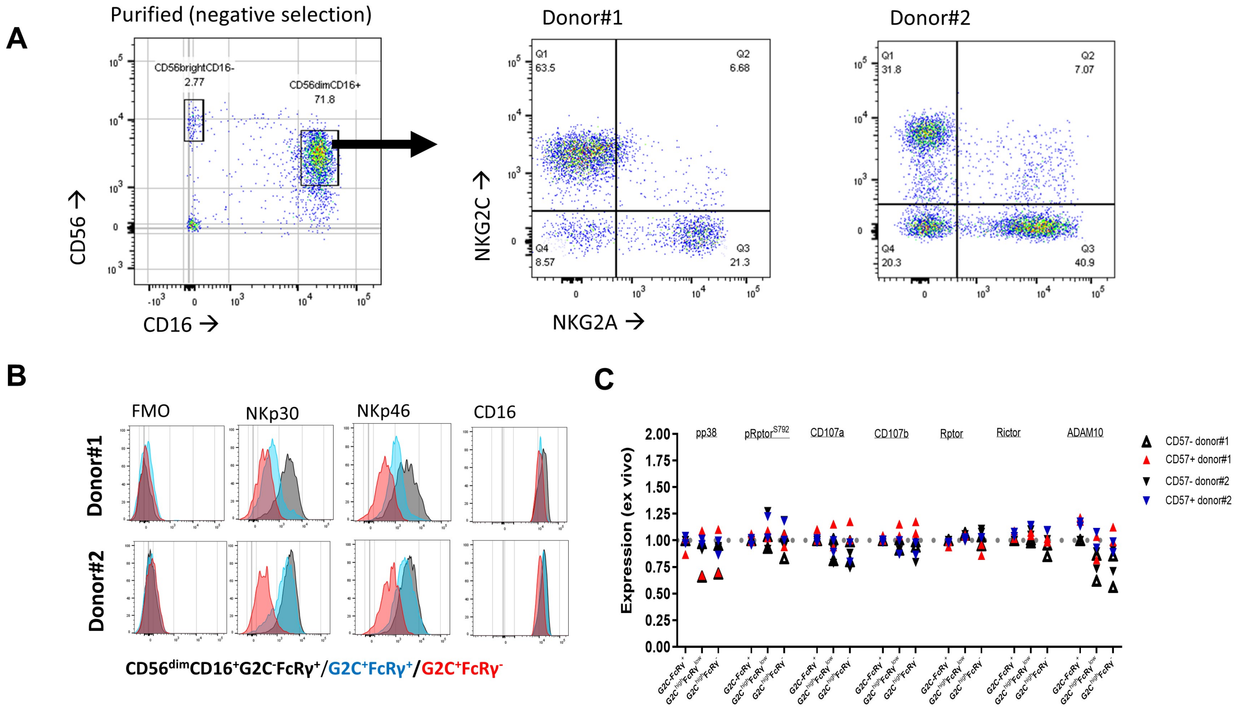


**Supplementary figure 2: ex vivo primary NK cells characterization and activation.**

**(A)** Representative dot plots showing gating of CD56$^{bright}$CD16$^{-}$, CD56$^{dim}$CD16$^{+}$, and the expression of NKG2C vs. NKG2A in CD56$^{dim}$CD16$^{+}$ cells from donor #1 and #2.

**(B)** Representative histograms for NKp30, NKp46, or CD16 expression, *ex vivo,* between the indicated NK cell sublets. (FMO = fluorescence minus one control)

**(C)** Intracellular pp38$^{Thr180/Tyr182}$, pRptor$^{S792}$, CD107a, CD107b, Rptor, or Rictor, and surface ADAM10 levels in *ex vivo* CD56$^{dim}$CD16$^{+}$ NK cells. Values were normalized to NKG2C$^{-}$FcRγ$^{+}$CD57$^{-}$ cells=1. Each measurement represents independent staining of an independent blood sample collected at an independent time point.

### Figure S3

**A**

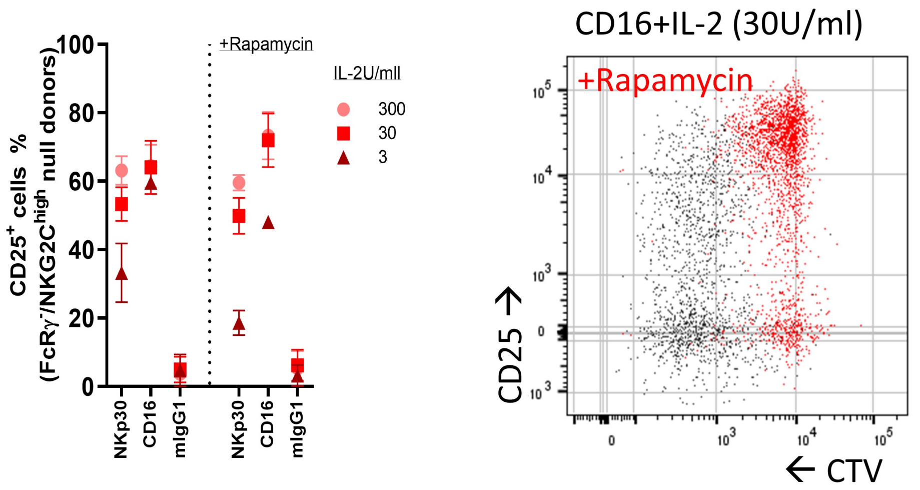


**B**

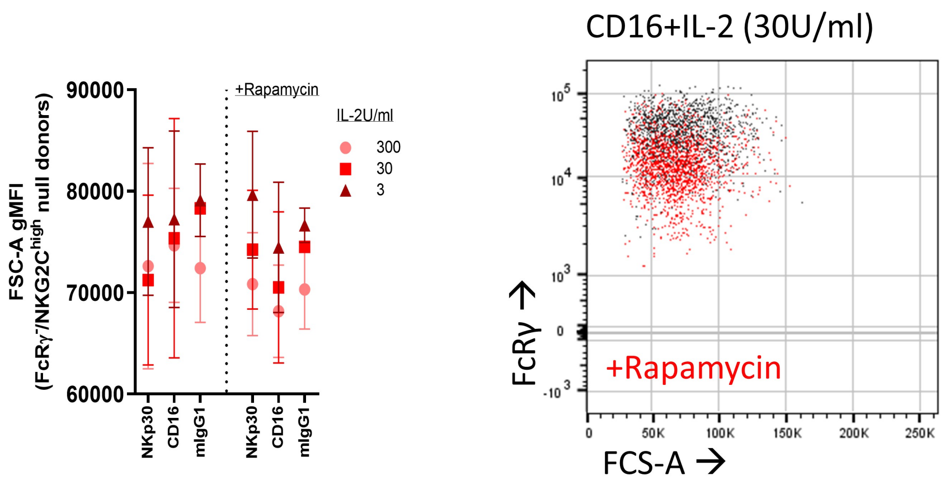


**Supplementary figure 3: rapamycin influence on CD25 upregulation and cell size during NK cell stimulation**

**(A)** Percentage of CD25$^{+}$ cells on total NK cells following 5 days of the indicated stimulation. Mean ± S.D, integrated results from two donors lacking NKG2C$^{high}$ or FcRγ$^{-}$ subsets (Donors #3 + #4). Right: representative dot plot of CD25 vs. CTV following anti-CD16 beads + IL-2 (30 U/ml) stimulation in the absence (black) or presence (red) of rapamycin (0.1 μM).

**(B)** Forward angle light scatter (FSC-A) gMFI levels of total NK cells following the indicated stimulation in the absence or presence of rapamycin (0.1 μM). Mean ± S.D. Integrated results from two donors lacking NKG2C$^{high}$ or FcRγ$^{-}$ subsets (Donors #3 + #4). . Right: representative dot plot of FcRγ expression vs. FSC-A following anti-CD16 beads + IL-2 (30 U/ml) stimulation in the absence (black) or presence (red) of rapamycin.
